# The oral protease inhibitor (PF-07321332) protects Syrian hamsters against infection with SARS-CoV-2 variants of concern

**DOI:** 10.1101/2021.11.04.467077

**Authors:** Rana Abdelnabi, Caroline S. Foo, Dirk Jochmans, Laura Vangeel, Steven De Jonghe, Patrick Augustijns, Raf Mols, Birgit Weynand, Thanaporn Wattanakul, Richard M. Hoglund, Joel Tarning, Charles E. Mowbray, Peter Sjö, Fanny Escudié, Ivan Scandale, Eric Chatelain, Johan Neyts

## Abstract

There is an urgent need for potent and selective antivirals against SARS-CoV-2. Pfizer developed PF-07321332 (PF-332), a potent inhibitor of the viral main protease (Mpro, 3CLpro) that can be dosed orally and that is in clinical development. We here report that PF-332 exerts equipotent *in vitro* activity against the four SARS-CoV-2 variants of concerns (VoC) and that it can completely arrest replication of the alpha variant in primary human airway epithelial cells grown at the air-liquid interface. Treatment of Syrian Golden hamsters with PF-332 (250 mg/kg, twice daily) completely protected the animals against intranasal infection with the beta (B.1.351) and delta (B.1.617.2) SARS-CoV-2 variants. Moreover, treatment of SARS-CoV-2 (B.1.617.2) infected animals with PF-332 completely prevented transmission to untreated co-housed sentinels.

## Introduction

There is an urgent need for potent and safe antiviral drugs for the treatment and prophylaxis of SARS-CoV-2 infections. Highly efficacious and safe viral proteases inhibitors have largely contributed to the effective treatment of infections with HIV and HCV. Coronaviruses have two proteases, the main protease Mpro (or 3CL protease) and the papain-like protease^1^. Mpro is a cysteine protease which cleaves the two polyproteins (pp1a and pp1ab) of SARS-CoV-2 at eleven different sites, resulting in the the various non-structural proteins, which are key for viral replication^2,3^. The substrate of Mpro is a distinct glutamine at the P1 site (Leu-Gln/Ser, Ala, Gly), no known human proteases recognize this cleavage site^4,5^. Mpro can thus be considered as a highly attractive drug target for the development of SARS-CoV-2 antivirals.

Pfizer reported recently that a SARS-CoV Mpro inhibitor which they developed in 2002 during the SARS-CoV-1 outbreak, is also effective against SARS-CoV-2^6^. Since the oral bioavailability of this compound is insufficient, an oral version, PF-07321332 (PF-332), was developed^7^. PF-332 potently inhibits the *in vitro* SARS-CoV-2 (USA_WA1/2020) replication as well as the replication of coronaviruses. The compound was also shown to exert antiviral activity in BALB/c mice infected with a mouse adapted SARS-CoV-2 variant (MA10)^7^.

We here report on the *in vitro* activity of PF-332 against different SARS-CoV-2 variants of concern (VoC) (including in human airway epithelial cultures) and on the efficacy of the drug in Syrian Golden hamsters (including in a transmission model) that had been infected with either the beta or the delta variant of SARS-CoV-2.

## Results

### *In vitro* antiviral activity of PF-332 against SARS-CoV-2 VoCs including in human airway epithelial cells (HAEC)

The *in vitro* antiviral activity of PF-332 against the four main SARS-CoV-2 VoC was first assessed in Vero E6 and A549 (overexpressing ACE2/TMPRSS2) cells, the EC_50_ values obtained were between 70 and 280 nM (**Table 1**). The antiviral effect of PF-332 was next assessed in primary human airway epithelial cell (HAEC) [that had been fully differentiated into an air-liquid (ALI) culture system] that were infected with the alpha variant (B.1.1.7). The assay has been previously validated for antiviral studies with SARS-CoV-2^8^. The parent nucleoside of remdesivir, GS-441524 (3 μM) was included as a reference inhibitor. When added to the culture medium at the basolateral site of the ALI’s 1h before infection (at the topical site) PF-332 (at 1 μM) completely inhibited viral replication for the entire duration of the experiment. At a concentration of 0.1 μM the inhibition was transient **(Fig 1)**.

**Table 1:**
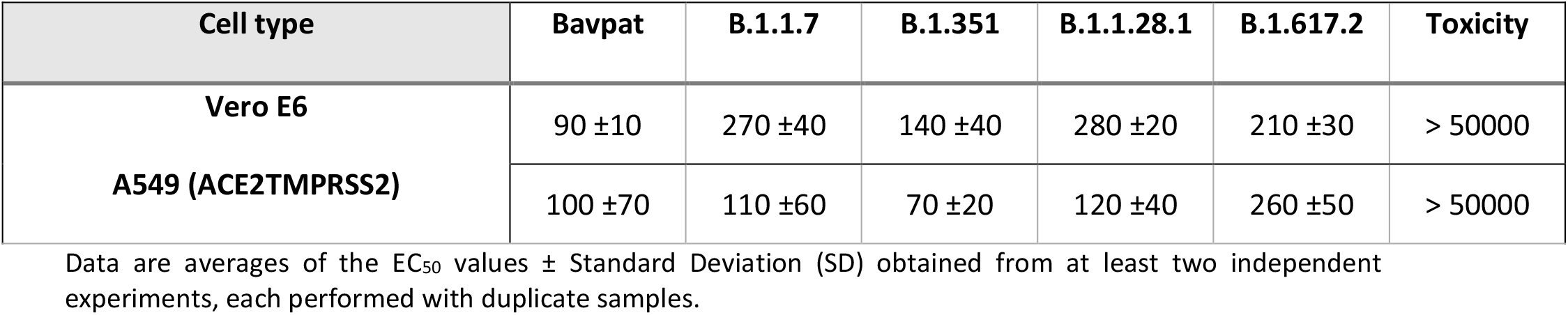
Antiviral activity (EC_50_ values in nM) of PF-332 against ancestral SARS-CoV-2 and variants in Vero E6-GFP cells and A549_ACE2TMPRSS2 cells.

| Cell type | Bavpat | B.1.1.7 | B.1.351 | B.1.1.28.1 | B.1.617.2 | Toxicity |
| --- | --- | --- | --- | --- | --- | --- |
| Vero E6 | 90 ±10 | 270 ±40 | 140 ±40 | 280 ±20 | 210 ±30 | > 50000 |
| A549 (ACE2TMPRSS2) | 100 ±70 | 110 ±60 | 70 ±20 | 120 ±40 | 260 ±50 | > 50000 |

Data are averages of the EC<sub>50</sub> values ± Standard Deviation (SD) obtained from at least two independent experiments, each performed with duplicate samples.

**Fig.1:**
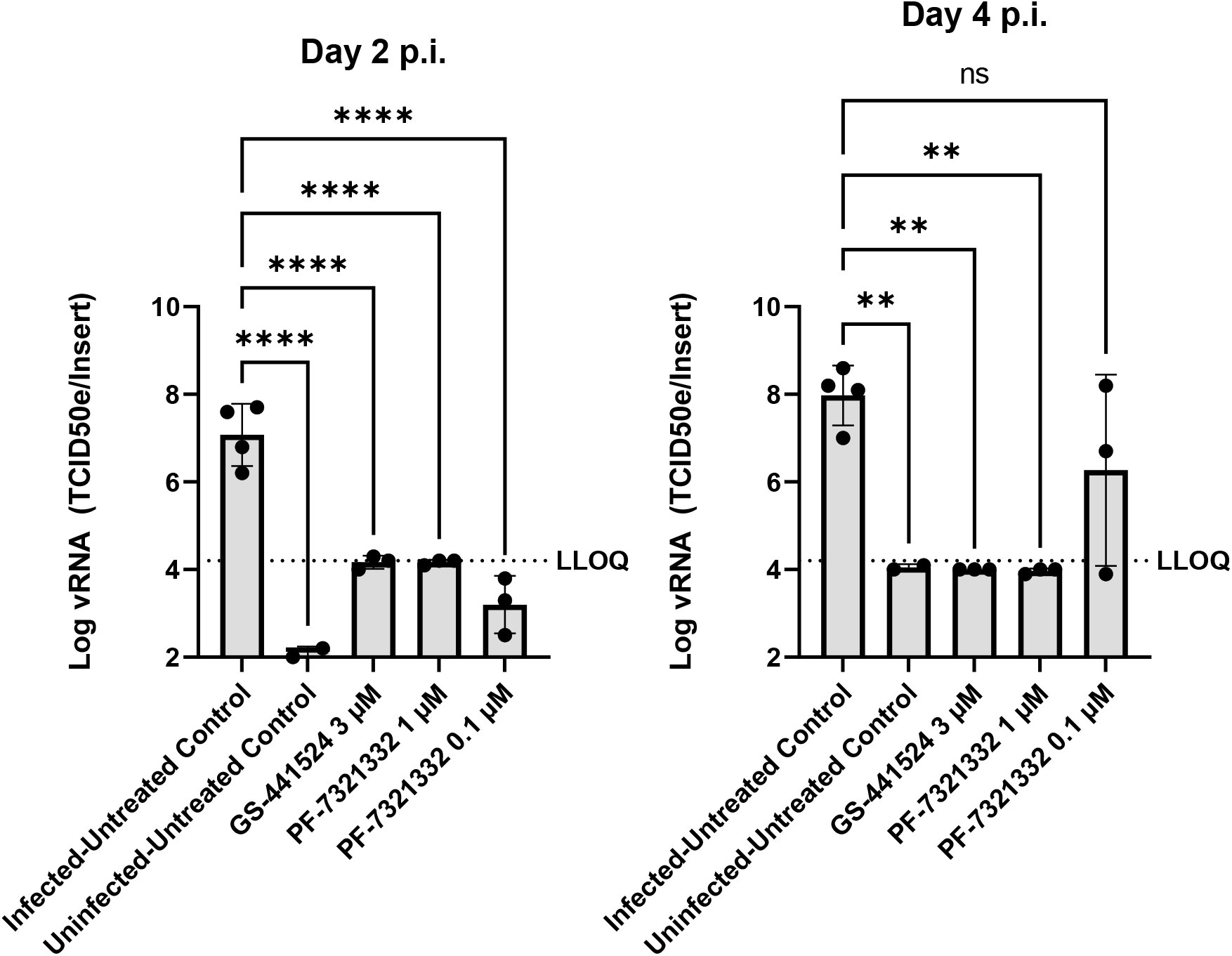
Antiviral activity of PF-332 in human Airway Epithelial cells infected with SARS-CoV-2 B1.1.7. Human airway epithelial cells, fully differentiated in an air-liquid culture system, were treated with compound at the basal site starting 1 h before infection with SARS-CoV-2 B.1.1.7 (alpha variant). Infection was done at the apical site. On day 2 and day 4 post viral RNA in culture supernatant was quantified. Each drug-treated condition is from 3 independent cultures. The infected/untreated control is from 4 independent cultures and the uninfected/untreated control from 2 independent cultures. Asterisks indicate the statistical significance between treated samples and the infected-untreated control. *p < 0.05, **p < 0.01, ***p < 0.001, ****p < 0.0001 (one-way ANOVA).

### PF-332 protects Syrian hamsters against challenge with the beta (B.1.351) SARS-CoV-2 VoC

Female hamsters (6 to 8 weeks) were intranasally infected with the SARS-CoV2 beta variant (lineage B.1.351) and were orally treated with PF-332 [either at 125 or 250 mg/kg/dose, twice daily (BID)] or the vehicle (i.e. the control group) for four consecutive days whereby treatment was initiated immediately before infection (**Fig. 2A**). Animals were euthanized at day four post-infection (pi). Treatment resulted in a dose-dependent reduction of viral RNA copies in lung tissue; i.e. 1.0 log_10_(P=0.010) and 5.9 log_10_ (P=0.002) reduction in respectively the 125 and 250 mg/kg, BID treatment groups (**Fig. 2B**). Likewise the 125 mg/kg BID dose resulted in a 1.1 log_10_ (P=0.015) reduction in lung infectious virus titers (**Fig. 2C**) and treatment with 250 mg/kg BID resulted in undetectable infectious virus levels in the lungs (4.8 log_10_ reduction, P=0.002) (**Fig. 2C**). Animals treated with the 125 mg/kg BID dose gained weight over the duration of the experiment whereas the vehicle group lost weight (**Fig. 2D**). Treatment also markedly improved virus-induced lung pathology, in particular in the 250 mg/kg BID dose whereby the lung pathology score was (in 5 out of 6 animals) comparable to the baseline score of untreated, non-infected hamsters (**Fig. 2E**).

**Fig.2.**
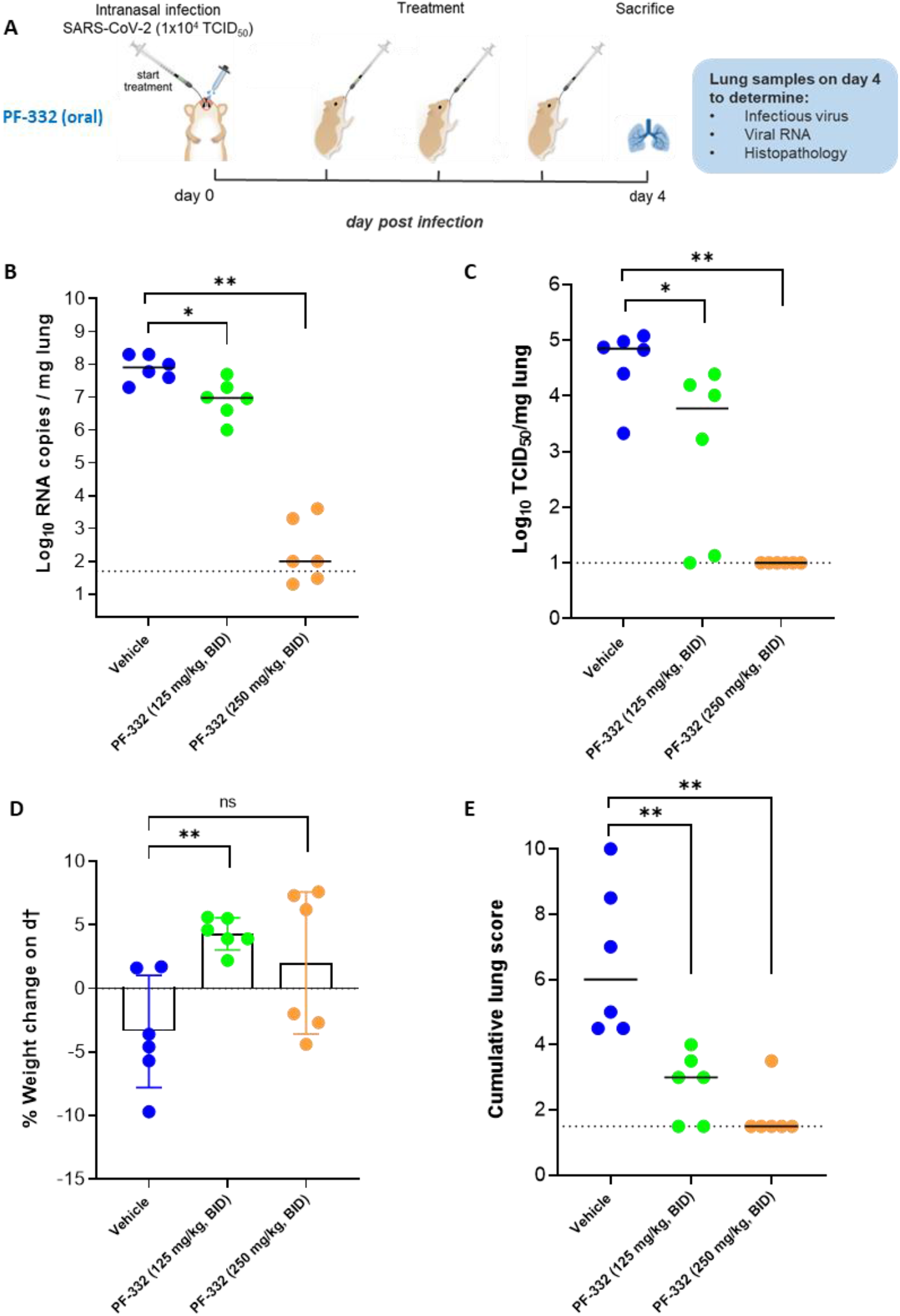
*In vivo* efficacy of PF-332 against Beta SARS-CoV-2 (B.1.351) variant in Syrian hamsters. (A) Design of the study. (B) Viral RNA levels in the lungs of control (vehicle-treated) and PF-332-treated (at 125 or 250 mg/kg, BID) SARS-CoV-2−infected hamsters at day 4 post-infection. Individual data and median values are presented and are expressed as log_10_ SARS-CoV-2 RNA copies per mg lung tissue. (C) Infectious viral loads in the lungs of control (vehicle-treated) and PF-332-treated SARS-CoV-2−infected hamsters at day 4 pi (expressed as log_10_ TCID_50_ per mg lung tissue). Individual data and median values are presented. (D) Weight change at day 4 pi in percentage, normalized to the body weight at the day of infection. Bars represent means ± SD. (E) Cumulative severity score from H&E stained slides of lungs from control (vehicle-treated) and PF-332-treated hamsters. Individual data and median values are presented and the dotted line represents the median score of untreated non-infected hamsters. Data were analysed with the Mann−Whitney U test. *P < 0.05, **P < 0.01, ns=non-significant (P<0.05). All data are from a single experiment with six animals per group. PF-332= PF-07321332.

### PF-332 protects Syrian hamsters against challenge with delta (B.1.617.2) SARS-CoV-2 VoC challenge and prevents viral transmission to untreated contact hamsters

We next studied whether treatment with PF-332 at 250 mg/kg BID also protects hamsters against challenge with the delta VoC (lineage B.1.617.2) and whether such treatment prevents transmission to untreated sentinels (**Fig. 3A**). To that end, index hamsters were intranasally infected and were treated from day 0 to day 2 with either vehicle or PF-332. On day 1 post-infection, which is immediately after administration of the morning dose to the index hamsters, each of the index hamsters was co-housed in a cage with contact/sentinel hamsters. The co-housing was continued until 2 days after start of contact (**Fig. 3A**). The index hamsters were then euthanized (day 3 pi) and the sentinels one day later. Treatment of index hamsters resulted in a 2.5 log_10_ (P=0.0022) reduction in viral RNA levels in the lungs and a 4.2 log_10_ reduction of infectious viral titres (P=0.0022), which is to undetectable levels (**Fig. 3B/C**). All sentinels that had been co-housed with vehicle-treated index hamsters had detectable viral RNA (**Fig. 3B**) and infectious virus loads in the lungs [ranging from 2×10^2^ to 6×10^4^ TCID_50_/mg lung tissue] (**Fig. 3C**). None of the sentinels that had been co-housed with PF-322-treated index hamsters had detectable infectious virus in the lungs (P=0.0022 compared to the contact of vehicle-treated group) (**Fig. 3C**).

**Fig.3.**
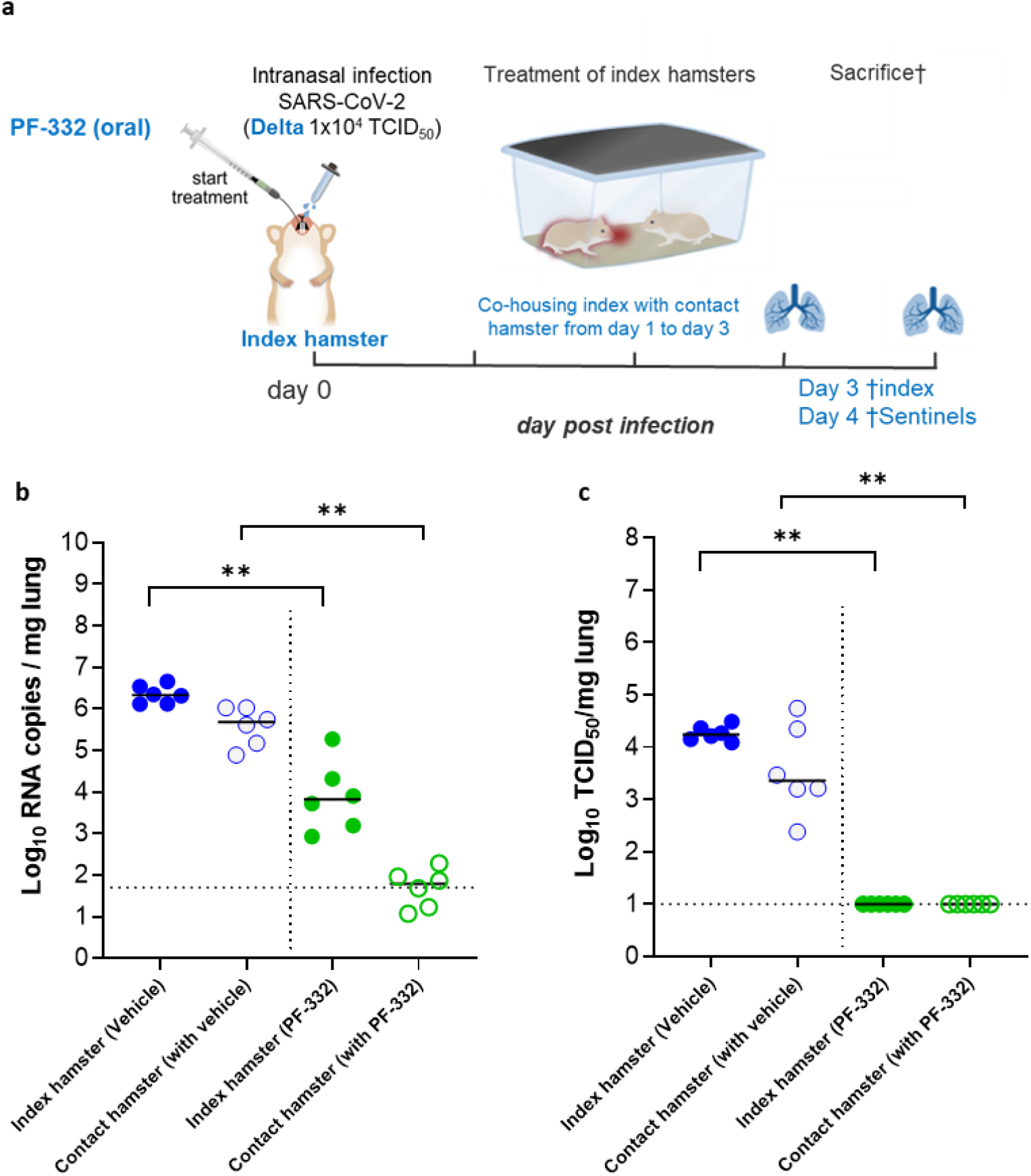
The effect of treatment with PF-332 on transmission of the Delta variant to untreated sentinel hamsters. (a) Design of the study. (b) Viral RNA levels in the lungs of control (vehicle-treated), PF-332-treated (250 mg/kg, BID) SARS-CoV-2−infected index hamsters (closed circles) and non-infected, non-treated contact hamsters (open circles) at day 3 and 4 post-infection (pi), respectively, are expressed as log_10_ SARS-CoV-2 RNA copies per mg lung tissue. Individual data and median values are presented. (c) Infectious viral loads in the lungs of control (vehicle-treated), PF-332-treated SARS-CoV-2−infected index hamsters and non-infected, non-treated contact hamsters at day 3 and 4 pi, respectively, are expressed as log_10_ TCID_50_ per mg lung tissue. Individual data and median values are presented. Data were analysed with the Mann−Whitney U test. **P < 0.01. PF-322=PF-07321332. The data are from a single experiment and with 6 animals per group.

### DMPK properties and pharmacokinetic analysis

Plasma protein binding was measured in mouse, hamster and human plasma. The average plasma protein binding expressed in percentage (± standard deviation) was respectively 74.0 (± 0.08), 37.9 (± 6.97), and 45.5 (± 1.5). The apparent intrinsic clearance (CL_int, app_) and the half-life measured (T_1/2_) measured following microsomal incubation were respectively for mouse, hamster, and human: CL_int, app_ = 73.6, 65.3, and 28.9 μl/min/mg; T_1/2_ = 23.5, 26.6, and 59.9 min.

Next a pharmacokinetic study was performed in uninfected hamsters. Animals received either a single oral dose of 50 mg/kg or 125 mg/kg (study 1) or a single oral dose of 125 mg/kg or 250 mg/kg (study 2) (cfr Methods). The pharmacokinetic properties were described by a two-compartment disposition model with first-order absorption; data are presented in **Supplementary Table 1**. A relatively fast terminal elimination half-life of 4-5 hours was calculated. Simulations revealed that (independent of the dosing scenario used) peak plasma concentrations of PF-332 are above the calculated *in vitro* EC_50_ (range from 70 nM – 260 nM) for any of the four variants in Vero E6 and A549 cells (**Supplementary Fig. S1 and S2, respectively**). Higher dosing resulted in longer duration of time above the target values associated with therapeutic efficacy. In addition, plasma levels of PF-332 in infected hamsters were determined at the time of sacrifice; in most animals an exposure of ∼100 nM was calculated (corresponding to 50 ng/ml) (**Table 2**) which is in the range of the EC_50_ determined in A549 cell cultures. The population pharmacokinetic analysis predicts plasma concentration at day four comparable to those obtained in infected animals. The average exposure was higher in the group treated with 250 mg/kg BID (**Table 2**), which resulted in these animals in a more pronounced antiviral effect than for the 125 mg/kg BID group.

**Table 2:** Comparison of PF-332 exposure in plasma at sacrifice (4 days post infection) with infectious virus titers in lungs.

| Treatment group | Hamster ID | TCID<sub>50</sub>/mg/lung | Plasma conc (nM) at d4† | Mean plasma conc (nM) at d4† |
| --- | --- | --- | --- | --- |
| Vehicle | 1 | 3·10<sup>4</sup> | 0 | 0 |
|  | 2 | 7·10<sup>4</sup> | 0 |  |
|  | 3 | 2·10<sup>3</sup> | 0 |  |
|  | 4 | 7·10<sup>4</sup> | 0 |  |
|  | 5 | 9·10<sup>4</sup> | 0 |  |
|  | 6 | 10<sup>5</sup> | 0 |  |
| PF-332 (125 mg/kg/BID) | 7 | 2·10<sup>3</sup> | 94 | 105 ± 35 |
|  | 8 | 0 | 122 |  |
|  | 9 | 2·10<sup>4</sup> | 39.0 |  |
|  | 10 | 10 | 134 |  |
|  | 11 | 2·10<sup>4</sup> | 118 |  |
|  | 12 | 10<sup>4</sup> | 125 |  |
| PF-332 (250 mg/kg/BID) | 13 | 0 | 122 | 275 ± 163 |
|  | 14 | 0 | 142 |  |
|  | 15 | 0 | 537 |  |
|  | 16 | 0 | 401 |  |
|  | 17 | 0 | 193 |  |
|  | 18 | 0 | 255 |  |

Average exposure in hamsters treated with 125 mg/kg, BID (105 ± 35 nM) is significantly lower (p<0.05) than exposure in hamsters treated with 250 mg/kg, BID (275 ± 163 nM), Unpaired Student T-test with equal distribution.

## Discussion

We here report that PF-332 (Pfizer) an oral SARS-CoV-2 Mpro inhibitor, protects Syrian Golden hamsters against challenge with both the beta and the delta VoC of SARS-CoV-2. To guide the hamster studies, we first assessed the *in vitro* antiviral activity of PF-332 against the four different VoCs. Nearly equipotent activity against the different variants was observed and this independent of the cell line used. PF-332 (at 1 μM) resulted also in full inhibition of the replication of the alpha VoC replication in primary human airways epithelial cells grown at the air-liquid interface which is a physiologically relevant model.

In hamsters infected with either the beta or the delta variant, a dose of 250 mg/kg BID of PF-332 reduced infectious virus titres in the lungs to undetectable levels. Half the dose (125 mg/kg BID) was highly effective in some but not all animals. Pharmacokinetic analysis corroborated these observations; indeed trough plasma PF-332 concentrations are maintained above the *in vitro* EC_50_ for the 250mg/kg BID dose but not consistently in animals that received the lower dose. Interestingly treatment of SARS-CoV-2 (delta) infected index hamsters completely prevented transmission to untreated co-housed sentinels. If such findings could be translated to the human situation, it may be assumed that treatment of infected patients will largely reduce a possible transmission to their contacts. Such strategy may have important consequences, for example in the context of a household when one or more members are infected and others not.

The antiviral effect of PF-332 in mice infected with a mouse-adapted SARS-CoV-2 strain was recently reported^7^. Oral BID dosing of PF-332 (at 300 and 1000 mg/kg) reduced titres of infectious virus in the lungs by respectively 1.4 and 1.9 log_10_ TCID_50_/ml as compared to the vehicle-treated group^7^. A marked reduction in lung histopathology was also observed in the lungs of PF-332-treated mice^7^. Obviously, even very high doses of PF-332 in mice do by far not achieve the pronounced antiviral potency as achieved in the hamster model (maximally 1.9 log_10_ reduction in infectious virus titres in the lungs of mice treated with 1000 mg/kg BID *versus* 4.2-4.8 log_10_ reduction in the lungs of hamsters treated with 250 mg/kg BID). This emphasizes the importance to assess the potential of drug candidates in various preclinical models. It should be noted that clinical isolates of the beta and the delta variant were used in the hamster study versus a mouse-adapted strain in the mouse studies.

Pharmacokinetic analysis revealed a relatively fast terminal elimination half-life of PF-332 of 4-5 hours. Co-administration with ritonavir, a potent CYP 3A4 inhibitor, might reduce the metabolism of PF-332 and increase the overall exposure, and/or reduce the dose needed. PF-332 is currently in clinical trials in combination with ritonavir (for example NCT05047601). Clearly, in the experiments presented here the exposures following administration of the highest dose were sufficient to achieve a pronounced antiviral effect. We will further explore whether the combined effect of PF-332 with ritonavir may result in a dose reduction of PF-332 in the hamster SARS-CoV-2 infection model.

Other direct antiviral drugs are in development for the treatment of COVID-19. As of today, only one antiviral drug has approved for the treatment of COVID-19, Veklury® (remdesivir). This injectable (intravenous dosing) treatment is exclusively indicated for hospitalized patients^9^. In a recent press release it was announced that early administration of Veklury® to high risk patients with mild to moderate symptoms reduces the likelihood of hospitalization with 87%. The active form remdesivir, which is the 5’-triphosphate of the parent nucleoside, is incorporated by the viral RNA dependent RNA polymerase (RdRp) into the growing RNA chain leading to chain termination^10^. An oral prodrug of the parent compound of remdesivir (i.e. GS-621763) has been developed and is results in promising results in preclinical models^11^. Molnupiravir (MK-4482, EIDD-2801), an oral prodrug of β-d-N4-hydroxycytidine, is an investigational oral antiviral medicine that was reported to result in a significant reduction of the risk of hospitalization or death during a phase 3 clinical trial when dosed to high risk patients with mild to moderate COVID19 [MOVe-OUT trial (NCT04575597). The 5’-triphosphate of β-d-N4-hydroxycytidine is incorporated by the RdRp in the viral RNA leading to lethal RNA mutagenesis^12^. We demonstrated that Molnupiravir is, alike PF-332 equipotent *in vitro* against the four VOC and is as well effective in the SARS-CoV-2 hamster infection model^13,14^. Yet another drug that is in clinical development, and authorized under emergency provisions, for the treatment of COVID-19 in a number of countries, including Japan, Russia, Turkey and India is favipiravir. A large cohort study with favipiravir is ongoing (PRINCIPLE trial). We reported earlier on the antiviral efficacy of favipiravir in the hamster model^15^. To achieve a maximal antiviral potency and to avoid at the same time the possible emergence of resistant variants, combination therapies may need to be developed. An oral Mpro inhibitor such as PF-332 may be an ideal candidate to be combined with a nucleoside analogue such as molnupiravir, favipiravir or an oral version of remdesvir.

In conclusion, we here demonstrate the antiviral efficacy of PF-332 against SARS-CoV-2 VoC *in vitro* and in the hamster infection model. Also infected animals treated with the drug, do no longer transmit the virus to untreated sentinels. Our data lend further support for the continued development of this drug.

## Methods

### Virus isolation and virus stocks

All virus-related work was conducted in the high-containment BSL3 facilities of the KU Leuven Rega Institute (3CAPS) under licenses AMV 30112018 SBB 219 2018 0892 and AMV 23102017 SBB 219 2017 0589 according to institutional guidelines.

The ancestral SARS-CoV-2 strain used for this study was derived from a German SARS-CoV-BavPat1/2020 isolate (hCoV-19/Germany/BY-ChVir-929/2020; EPI_ISL_406862; 2020-01-28, kindly provided by C. Drosten, Charité, Berlin, Germany). The four SARS-CoV-2 Variants of Concern (VoC) used in this study were Alpha B.1.1.7 (derived from hCoV-19/Belgium/rega-12211513/2020; EPI_ISL_791333, 2020-12-21) ^16^, Beta B.1.351 (derived from hCoV-19/Belgium/rega-1920/2021; EPI_ISL_896474, 2021-01-11) ^16^, Gamma P.1 (EPI_ISL_1091366; 2021-03-08) and Delta B.1.617.2 (derived from hCoV-19/Belgium/rega-7214/2021; EPI_ISL_2425097; 2021-04-20). The four variants were originally isolated in-house from nasopharyngeal swabs taken from travellers returning to Belgium (baseline surveillance) and were subjected to sequencing on a MinION platform (Oxford Nanopore) directly from the nasopharyngeal swabs ^16^. Virus stocks were then grown on Vero E6 cells in (DMEM 2% FBS medium) and passaged two times. Median tissue culture infectious doses (TCID_50_) was defined by end-point titration as described before ^16^.

### Compounds

PF-07321332 was synthesized at TCG Lifesciences (India) and Wuxi (USA). GS-441524 (the parent nucleoside analogue of remdesivir) was purchased from Carbosynth (United Kingdom). For *in vitro* assays, compounds were dissolved in analytical grade dimethyl sulfoxide (DMSO) to 10mM stock solution. For *in vivo* studies, the Beta variant efficacy study was performed with PF-07321332 (TCG Lifesciences) that was formulated as 40 mg/ml in a vehicle containing 40% PEG400 (Sigma) in sterile distilled water. PF-07321332 (from Wuxi) was used only in the delta variant hamster study and it was formulated as 40 mg/ml in a vehicle containing 60% PEG400 (Sigma) in sterile distilled water. In these studies, the crystalline form of the active product ingredient (API) was characterized by differential scanning calorimetry (DSC) (data not shown). X-ray powder diffraction (XRPD) would be however required to fully characterize the crystalline form of each API used in *in vivo* studies.

### SARS-CoV-2 *in vitro* antiviral assays

The assay using Vero E6 cells was derived from a previously established SARS-CoV assay ^17^. In this assay, fluorescence of Vero E6-eGFP cells (provided by Dr. K. Andries J&JPRD; Beerse, Belgium) declines after infection with SARS-CoV-2 due to virus-induced cytopathogenic effect. In the presence of an antiviral compound, the cytopathogenicity is inhibited and the fluorescent signal maintained. Vero E6 cells were maintained in Dulbecco’s modified Eagle’s medium (DMEM; Gibco cat no 41965-039) supplemented with heat-inactivated 10% v/v foetal calf serum (FCS; HyClone) and 500 μg/mL Geneticin (Gibco cat no 10131-0275) and kept under 5% CO2 at 37°C.

The test compounds were serially diluted in assay medium (DMEM supplemented with 2% v/v FCS). Diluted compounds were then mixed with Vero E6-eGFP cells corresponding to a final density of 25,000 cells/well in 96-well blackview plates (Greiner Bio-One, Vilvoorde, Belgium; Catalog 655090). The next day, cells were infected with the SARS-CoV-2 at a final MOI of approximately 0.05 TCID_50_/cell. Final dilution of the different strains was adapted in order to obtain a similar MOI between all variants of interest. The plates were incubated in a humidified incubator at 37°C and 5% CO2. At 4 days post-infection (pi), the wells were examined for eGFP expression using an argon laser-scanning microscope. The microscope settings were excitation at 488 nm and emission at 510 nm and the fluorescence images of the wells were converted into signal values. Toxicity of compounds in the absence of virus was evaluated in a standard MTS-assay as described previously ^18^.

A549-Dual™ hACE2-TMPRSS2 cells obtained by Invitrogen (Cat. a549d-cov2r) were cultured in DMEM 10% FCS (Hyclone) supplemented with 10 μg/ml blasticidin (Invivigen, ant-bl-05), 100 μg/ml hygromycin (Invivogen, ant-hg-1), 0.5 μg/ml puromycin (Invivogen, ant-pr-1) and 100 μg/ml zeocin (Invivogen, ant-zn-05). For antiviral assay, cells were seeded in assay medium (DMEM 2%) at a density of 15,000 cells/well. One day after, compound was serially diluted in assay medium (DMEM supplemented with 2% v/v FCS) and cells were infected with their respective SARS-CoV-2 strain at a MOI of approximately 0.05. The MOI was kept comparable for the variant strains in the different experiments. On day 4 pi., differences in cell viability caused by virus-induced CPE or by compound-specific side effects were analyzed using MTS as described previously ^18^.

The results of *in vitro* antiviral experiments were expressed as EC_50_ values defined as the concentration of compound achieving 50% inhibition of the virus-reduced eGFP signals as compared to the untreated virus-infected control cells.

### Human Airway Epithelial model

#### Viral infection

Tracheal HAEC (catalogue no. EP01MD) from healthy donors were provided by Epithelix company (Geneva, Switzerland) in an air-liquid interphase set-up and treated as described elsewhere ^8^. On day 0 of the experiment, the H(s)AEC were first pre-treated with basal medium containing compounds, followed by infection with SARS-CoV-2_B.1.1.7 at 5×10^2 TCID_50_/insert virus input at the apical side for 1.5 hours. The first apical wash with MucilAir medium was collected at day 1 pi. Every other day from day 0, subsequent apical washes were collected, whereas compound-containing medium in the basolateral side of the H(s)AEC culture was refreshed. Wash fluid was stored at -80°°C until analysis by RT-qPCR.

#### RNA extraction and quantitative reverse transcription-PCR (RT-qPCR)

Viral RNA in the apical wash was isolated using the Cells-to-cDNA II cell lysis buffer kit (Thermo Fisher Scientific, catalogue no. AM8723). Briefly, 5 μL wash fluid was added in 50 μL lysis buffer, incubated at room temperature (RT) for 10 min and then at 75°C for 15 min. 150 μL nuclease-free water was additionally added to the mixture prior to RT-qPCR. Together with the samples, a ten-fold serial dilution of the corresponding virus stock was extracted to later generate a standard curve for the RT-qPCR. Based on this standard curve the amount of viral RNA can be expressed as median Tissue Culture infective dose (TCID_50_) equivalents per insert (TCID_50_eq/insert), and the lowest point of the linear part of the standard curve (highest Ct value) determines the lower limit of quantification (LLOQ). The RT-qPCR was performed using iTaq universal probes one-step kit (Bio-Rad, catalogue no. 1725141), and a commercial mix of primers for N gene, manufactured at IDT Technologies (catalogue no. 10006606). The reaction (final volume: 20 μL) consisted of 10 μL one-step reaction mix 2x, 0.5 μL reverse transcriptase, 1.5 μL of primers and probes mix, 4 μL nuclease-free water, and 4 μL viral RNA. The RT-qPCR was executed on a Lightcycler 96 thermocycler (Roche), starting at 50°C for 15 min and 95°C for 2 min, followed by 45 cycles of 3 sec at 95°C and 30 sec at 55°C.

### SARS-CoV-2 infection model in hamsters

The hamster infection model of SARS-CoV-2 has been described before ^15,19^. Female Syrian hamsters (Mesocricetus auratus) were purchased from Janvier Laboratories and kept per two in individually ventilated isolator cages (IsoCage N Bio-containment System, Tecniplast) at 21°C, 55% humidity and 12:12 day/night cycles. Housing conditions and experimental procedures were approved by the ethics committee of animal experimentation of KU Leuven (license P065-2020). For infection, female hamsters of 6-8 weeks old were anesthetized with ketamine/xylazine/atropine and inoculated intranasally with 50 μL containing 10^4^ TCID_50_ of SARS-CoV-2 Beta variant B.1.351 (day 0). On day 4 pi, animals were euthanized for sampling of the lungs and further analysis by i.p. injection of 500 μL Dolethal (200 mg/mL sodium pentobarbital, Vétoquinol SA). All caretakers and technicians were blinded to group allocation in the animal facility.

#### Treatment regimen (Beta variant study)

Hamsters (as n=6 per group) were treated by oral gavage with either the vehicle or PF-332 at 125 or 250 mg/kg/dose twice daily starting from D0, just before the infection with the Beta variant. All the treatments continued until day 3 pi. Hamsters were monitored for appearance, behavior and weight. At day 4 pi, hamsters were euthanized by i.p. injection of 500 μL Dolethal (200mg/mL sodium pentobarbital, Vétoquinol SA). Lungs were collected and viral RNA and infectious virus were quantified by RT-qPCR and end-point virus titration, respectively as described before ^16^.

#### Efficacy-transmission study (Delta variant study)

Two groups of index hamsters were infected with Delta variant and treated with either vehicle or PF-332 at 250 mg/kg/dose twice daily starting from D0. On day1 pi (just after the morning dose), each index hamster was co-housed with a contact hamster (non-infected, non-treated hamsters) in one cage and the co-housing continued until day 3 pi The treatment of index hamsters was continued until day2 pi. At day 3 pi, all the index hamsters were euthanized whereas all the contact hamsters were euthanized the day after (i.e. day 4 pi of index) as mentioned before and lungs were collected to assess viral loads.

#### Histology

For histological examination, the lungs were fixed overnight in 4% formaldehyde and embedded in paraffin. Tissue sections (5 μm) were analyzed after staining with hematoxylin and eosin and scored blindly for lung damage by an expert pathologist. The scored parameters, to which a cumulative score of 1 to 10 was attributed, were the following: congestion, intra-alveolar hemorrhagic, intra-alveolar edema, apoptotic bodies in bronchus wall, necrotizing bronchiolitis, perivascular edema, bronchopneumonia, perivascular inflammation, peribronchial inflammation and vasculitis.

#### Sample size justification

For *in vivo* antiviral efficacy, we want to detect at least 1 log_10_ reduction in viral RNA levels in treated subjects compared to the untreated, infected control group. Group size was calculated on the independent t-test with an effect size of 2.0 and a power of 80% (effect size = deltamean/SD = 1 log_10_ decrease in viral RNA/0.5 log_10_), resulting in 5-6 animals/group. Sample sizes maximized considering limits in BSL3 housing capacity, numbers of animals that can be handled under BSL3 conditions, and availability of compounds.

#### Statistics

The number of animals and independent experiments that were performed is indicated in the legends to figures. The analysis of histopathology was done blindly. All statistical analyses were performed using GraphPad Prism 9 software (GraphPad, San Diego, CA, USA). Statistical significance was determined using the non-parametric Mann Whitney U-test or student T-test. P-values of <0.05 were considered significant.

### *In vitro* drug metabolism and pharmacokinetics (DMPK)

#### Protein binding

Plasma protein binding was measured by rapid equilibrium dialysis (RED) method to determine the free fraction and the unbound percentage of PF-332 for various species. An equilibrium dialysis was conducted in duplicate for each sample. 200μl of plasma spiked with PF-332 were added in the plasma chamber and 350 μl of PBS pH = 7.4 were added in the buffer chamber. The dialysis block was then incubated at 37°C for 6 hours with constant shaking at 400rpm (Thermomixer comfort, Eppendorf). After 6 hours, aliquots of the plasma and the buffer chambers were collected, spiked to obtain a matching homogeneous matrix, and quantified by LC-MS / MS (API-4000 MS, AB Sciex Instruments, coupled with Shimadzu LC-20ADvp Prominence Liquid Chromatography and CTC Analytics HTS PAL autosampler).

The percentages of the free concentration were respectively calculated using following formulas:

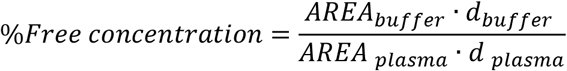

Where ‘d’ represents the dilution factor and ‘AREA’ the chromatographic peak area of the analyte. As the percentage of the coefficient of variation (CV) of the Internal standard is below 10%, the analyte area is considered for calculation instead of internal standard-normalised peak area.

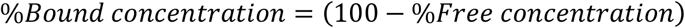

#### Microsomal metabolic stability

Mouse liver microsomes (CD-1 male strain) were purchased from GIBCO. Hamster (Syrian female strain) and human liver microsomes were purchased from Xenotech. 1 ml of liver microsomal (LM) suspension at 20 mg/ml was mixed with 19 ml of 100 mM phosphate buffer. The latter is a titer solution containing 1 (M) KH_2_PO_4_ and 1 (M) K_2_HPO_4_ diluted in 10-fold distilled water (30 ml buffer + 270ml of water) to obtain 100mM phosphate buffer with an adjusted pH at 7.40 ± 0.02. A solution of NADPH Regeneration System (NRS) was prepared using 13 mM NADP, 33 mM Glucose-6-phosphate, 33 mM MgCl_2_, and 4 U/ml buffer solution of Glucose-6-phosphate dehydrogenase.

All plastic materials including tips are incubated at 37°C overnight. The LM suspension and the NRS solution were incubated at 37°C for ∼15min before use. 48μl of buffer was added to the wells of the blank plate. 40 μl of the compound at 1 uM was added to the working plates, 8μl of NRS solution was added in the 0, 5, 10, 20, 30, and 60min plates. The reaction is then initiated by adding 32μl of 1 mg/mL of LM suspension to each plate. The reaction is terminated by adding 240μl ice-cold acetonitrile at the designated time points. At T = 0, the acetonitrile is added before the LM solution.

The plates are centrifuged (3500rpm, 20min and 15°C); 110μl of distilled water are then added to 110μl of the supernatant and analyzed using an LC-MS/MS (API-4000 MS, AB Sciex Instruments, coupled with Shimadzu LC-20ADvp Prominence Liquid Chromatography and CTC Analytics HTS PAL autosampler). The apparent *in vitro* intrinsic clearance is then calculated using the following formulas

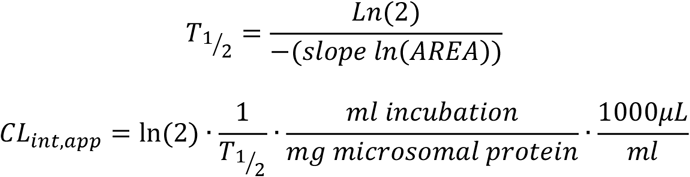

Where T_1/2_ is the half-life and CL_int, app_ the apparent *in vitro* intrinsic clearance.

### Population pharmacokinetic analysis

The pharmacokinetic samples used for population pharmacokinetic analysis were collected in female Syrian hamsters from two satellite studies (study 1 and study 2) with uninfected hamsters sourced from Hylasco Biotechnology – a Charles River licensee) and LIVEON BIOLABS (Bangalore). In both studies, the pharmacokinetic samples were collected at 0, 0.5, 1, 3, 6, 12, 24h after the dose was given orally. In study 1, the samples were collected from 8 animals that received a single oral dose of 50 mg/kg (n=4) and 125 mg/kg (n=4) formulated as a homogenous suspension in 0.5% methylcellulose (w/v) in water. In study 2, samples were collected from 6 animals that received a single oral dose of 125 mg/kg (n=3) and 250 mg/kg (n=3) formulated as a homogenous suspension of 40% PEG 400, 60% MilliQ water. The animals were fasted 12h before the dose and until 4h after dose.

At each time point, approximately 80 ul of blood from the female Syrian hamster were collected by pricking lateral saphenous vein with a 26G needle. All samples were transferred into commercial Heparin/K2-EDTA tubes, placed on ice until processed for plasma extraction by centrifugation and stored at -20°C before analysis. A LC-MS/MS platform (API 4500/4000 MS, Shimadzu LC and CTC autosampler) was used to determine PF-332 concentrations with a reverse-phase LC-MS/MS method developed with an internal standard. The mobile phase was a gradient of 0.1% formic acid (FA) in water the column was an Kinetex Biphenyl, 2.1*30mm.

The data from the two studies were modelled simultaneously using nonlinear mixed-effects approach in NONMEM, v7.4 (Icon Development Solution, Ellicott City, MD)^20,21^. One-, two-, and three-compartment models were evaluated to describe the structural disposition model of PF-332. First-order and zero-order absorption models were evaluated to describe drug absorption. Relative bioavailability was fixed to unity and added into the model by estimating the inter-individual variability. In order to improve the translational aspect of the model, body weight was implemented on all clearance and volume parameters using a fixed allometric function with an exponent of 0.75 and 1 for clearance and volume, respectively. Differences in the pharmacokinetic properties between the two satellite studies were expected due to the manufacturing process of PF-332, essentially resulting in different formulations of PF-332. These differences were evaluated as a categorical covariate on all pharmacokinetic parameters. Additionally, dose was also evaluated as a continuous linear covariate in the model. The covariate investigation was performed using a stepwise forward inclusion (P=0.05, ΔOFV = –3.84), followed by a stepwise backward elimination (P=0.001, ΔOFV= –10.83) procedure. Parameter precision was obtained using a bootstrapping approach with 1,000 re-sampled datasets stratified by study. Predictive performance was evaluated with simulation-based diagnostics (**Supplementary Fig. S3**).

The pharmacokinetic parameters from the population pharmacokinetic model were used to simulate 1,000 concentration-time profiles for each dosing scenario: 50 mg/kg single dose, 125 mg/kg single dose, 125 mg/kg twice daily for 4 consecutive days, and 250 mg/kg twice daily for 4 consecutive days. All the simulations were performed using Simulx2020R1 (Lixoft, Batiment D, Antony, France). The simulation results were overlaid with the *in vitro* EC_50_ reported in Vero E6 cells and A549-ACE2TMPRSS2 cells. These EC_50_ values were corrected for a plasma protein binding of 37.8% reported in hamsters, in order to compare the simulated total drug concentration-time profiles from the developed pharmacokinetic model with expected therapeutic concentrations.

### Compound exposure determination in infected hamsters

Plasma samples were obtained at end point of the study (during sacrifice) and inactivated by 1h UV exposure. Samples were analysed using RP-UPLC with tandem mass spectrometry detection (Acquity H-class UPLC, Waters, Milford, MA, USA and Xevo TQ-S micro Waters, Milford, MA, USA). In brief, plasma samples were analysed for PF-332 after protein precipitation with methanol (containing the internal standard propranolol at 10nM). Chromatographic separation was performed using a Kinetex XB -C18 column (2.6 μm, 2.1 × 50 mm; Phenomenex, Utrecht, the Netherlands) held at 40°C. Methanol (solvent A) and 0.05% formic acid in water (solvent B) were used as eluents at 500 μL/min. Gradient elution was performed as follows: 0-0.5 min, 5% solvent A; 0.5-0.6 min, 5→84% solvent A; 0.6-1.7 min, 84% solvent A; 1.7-1.8 min 84→95% solvent A; 1.8-2.8 min, 95% solvent A; 2.8→3.0 min, 95-5% solvent A; 3.0-4.5 min, 5% solvent A to re-equilibrate the column prior to the next injection. Propranolol and PF-332 eluted at 1.45 and 1.62 min, respectively.

MS/MS was carried out with a HESI source in the positive ionization mode. Detection and quantification were performed using selected reaction monitoring (SRM) for the transitions m/z 260.20→116.10 for propranolol and 500.30→319.20 for PF-332. Calibration curves were made on the day of analysis by serial dilution in plasma. The method was proven to be linear, accurate and precise over the range of 0.49-1000 nM.

## Ethics

Housing conditions and experimental procedures were done with the approval and under the guidelines of the ethics committee of animal experimentation of KU Leuven (license P065-2020).

## Data Availability

All of the data generated or analysed during this study are included in this published article.

## Acknowledgments

We thank Carolien De Keyzer, Lindsey Bervoets, Thibault Francken, Birgit Voeten, Dagmar Buyst, Niels Cremers for excellent technical assistance. We are grateful to Piet Maes for kindly providing the SARS-CoV-2 strain used in this study. We thank Jef Arnout and Annelies Sterckx (KU Leuven Faculty of Medicine, Biomedical Sciences Group Management) and Animalia and Biosafety Departments of KU Leuven for facilitating the studies.

## Author Contribution

All authors read and approved the final version of the manuscript. Study concept and design /conceptualization: RA, EC, FE, IS, DJ, JN. Project administration / Coordination: RA, CSF, EC, FE, IS, LV, DJ, JN. Data analysis: RA, CSF, TW, RH, JT, RM, PA, BW, DJ. Method: RA, CSF, LV, RM, BW, DJ. Writing— review and editing: RA, TW, RMH, JT, EC, IS, FE, CEM, PS, CSF, RM, PA, DJ, JN. Access to essential reagents: SDJ. Funding acquisition: EC, CEM, IS, FE, PS, LV, DJ, JN.

## Supplementary

**Fig.S1.**
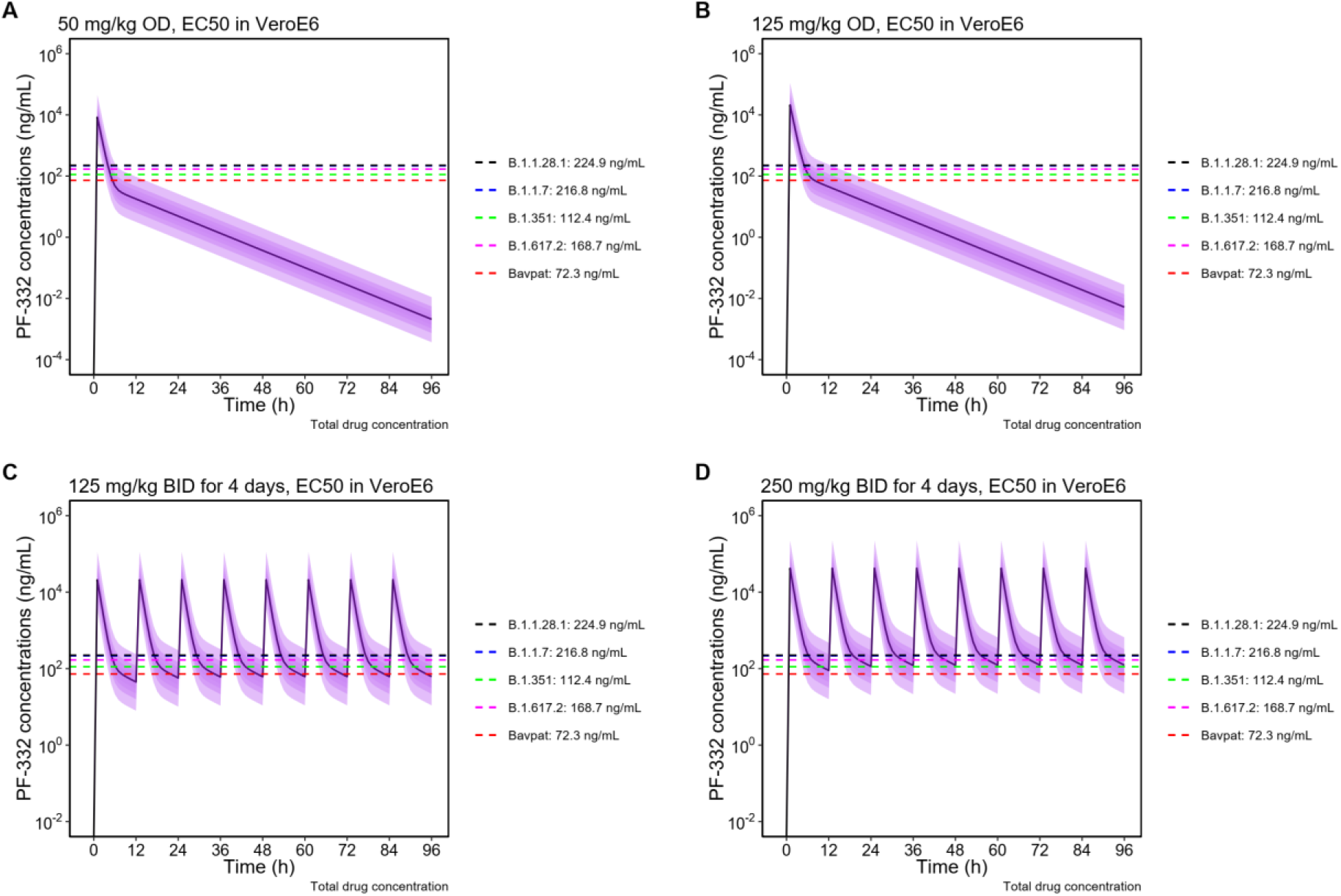
The simulated concentration time profile of PF-332 with different dosing scenarios (n=1000 for each scenario) and compared with EC_50_ in Vero E6 cell line: (A) 50 mg/kg once daily, (B) 125 mg/kg once daily, (C) 125 mg/kg twice daily for 4 days, (D) 250 mg/kg twice daily for 4 days. The horizontal lines represent the EC_50_ values reported in Vero E6 cell which is corrected for 37.8% plasma protein binding in hamsters.

**Fig.S2.**
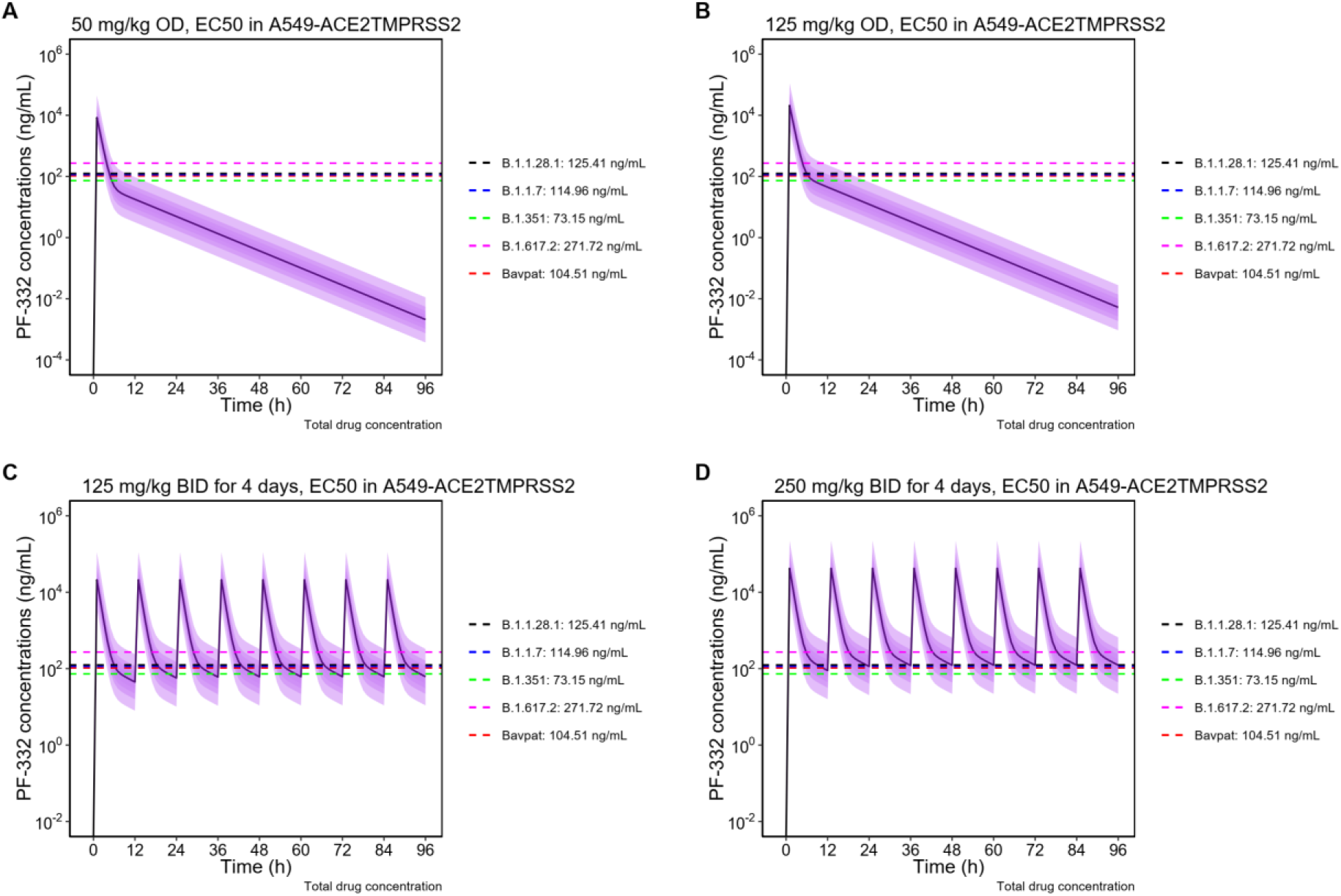
The simulated concentration time profile of PF-332 with different dosing scenarios (n=1000 for each scenario) and compared with EC_50_ values obtained in A549 cell line: (A) 50 mg/kg once daily, (B) 125 mg/kg once daily, (C) 125 mg/kg twice daily for 4 days, (D) 250 mg/kg twice daily for 4 days. The horizontal lines represent the EC_50_ values reported in A549-ACE2TMPRSS2 cell which is corrected for 37.8% plasma protein binding in hamsters.

**Fig.S3.**
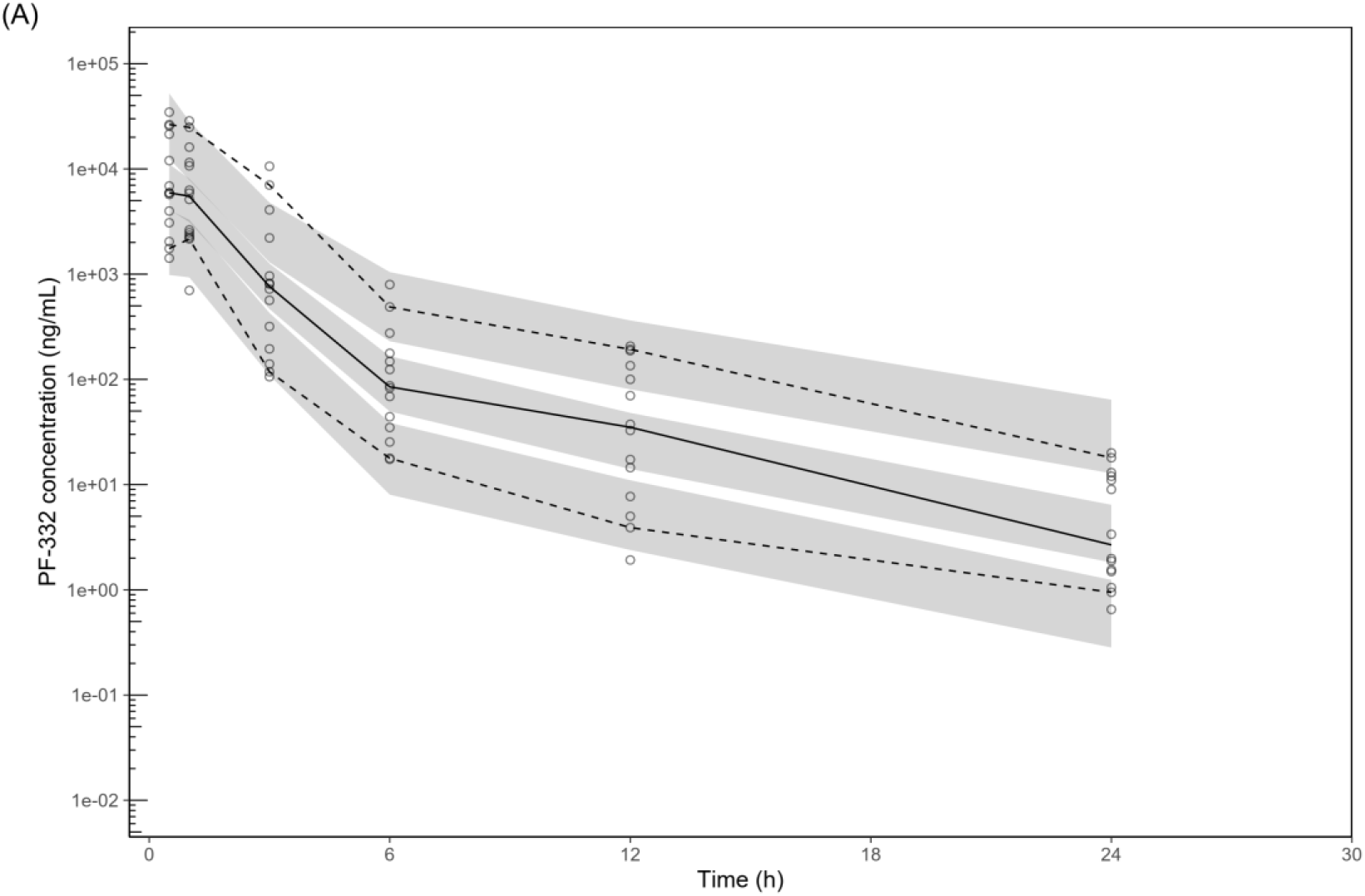
Visual predictive check of the final population pharmacokinetic model of PF-332 in hamster. The open circles represent the observed PF-332 concentrations. Solid black lines represent the 50th percentiles of the observations, and dashed black lines represent the 5th and 95th percentiles of the observations. The shaded areas represent the 95% confidence intervals of each simulated percentile (n =1,000).

**Table S1.** Population pharmacokinetics parameters of PF-332 in hamsters.

| Parameter | Population estimates <sup>a</sup><br>(%RSE) <sup>b</sup> | 95%CI<sup>b</sup> | IIV <sup>a</sup> [%CV]<br>(%RSE) <sup>b</sup> | 95%CI<sup>b</sup> |
| --- | --- | --- | --- | --- |
| F | 1 (fixed) | - | - | - |
| CL/F (ml/min) | 9.83 (24.5) | 5.88-15.5 | 5.40 (69.9) | 0.0541 –<br>15.8 |
| V/F (ml) | 75.0 (89.5) | 14.5-364 | - | - |
| Q/F (ml/min) | 0.292 (56.5) | 0.0952-0.760 | - | - |
| VP/F (ml) | 93.5 (55.7) | 31.8-250 | - | - |
| k<sub>a</sub> (min<sup>-1</sup>) | 0.0215 (14.2) | 0.0197-0.0303 | - | - |
| Covariate effect compared to study 1 |  |  |  |  |
| Study 2-CL (%<br>lower) | 89.3 (6.15) | 75.1 – 95.2 | - | - |
| Study 2-F (% lower) | 95.8 (4.65) | 83.7 – 98.9 | - | - |
| Ⓜ | 0.482 (7.78) | 0.303-0.562 | - | - |

<sup>a</sup> Population mean values, inter-individual variability (IIV) were estimated by NONMEM. The coefficient of variation (%CV) for IIV were calculated as $100 \times \sqrt{\exp(\text{estimate}) - 1}$ .
<sup>b</sup> Relative standard error (%RSE) was calculated as $100 \times \left( \frac{SD}{\text{Mean value}} \right)$ from the non-parametric bootstrap results (n=1,000). The 95% confidence interval (95%CI) is presented as the 2.5 to 97.5 percentiles of bootstrap estimates.

